# Counterfactual aging in individual digital twin brains uncovers a latent dimension of resilience to aging

**DOI:** 10.64898/2026.11.18.767485

**Authors:** Yuta Takahashi, Hayato Idei, Hiroaki Tomita, Yuichi Yamashita

## Abstract

Cognitive reserve concerns why the same aging causes less decline in some individuals than others, a question that digital twin brains can address by aging each individual in silico. Here we present Brain-VRNN, a connectome-constrained variational recurrent neural network trained on longitudinal multitask functional MRI. Its regenerated brain signals reproduced four functional aging markers of unseen individuals (r = 0.93–0.96) and identified each person (accuracy 0.98). Without cognitive scores, its global latent learned a trait dimension stable within each person, which helped predict future cognition. Crucially, when every individual underwent the same counterfactual aging, those with a lower trait value declined more, yet recovered up to 1.6 times as much with a reduction in body mass index, a modifiable dementia risk factor. Aging single regions showed that different markers decline through different regions, with a lateral temporal–angular region shared by all. Brain-VRNN brings within-person simulation to cognitive reserve.

## INTRODUCTION

Why the same aging leads to different degrees of cognitive decline in different individuals has long been studied under the concepts of cognitive reserve and, more broadly, resilience to aging^1–3^. Neuroimaging studies have described this variability in terms of compensation and dedifferentiation^4,5^, suggesting that part of the answer lies in how the contribution of each region, whose activity propagates to others through their connections, is reorganized with age. This reorganization has been examined primarily through correlational analyses of observational data, which cannot compare the same individual with and without a change in age or in a modifiable condition. Digital twin brains, mathematical models that generate the signals of an individual’s brain and simulate how these signals respond to an intervention, are attracting growing interest in cognitive aging and dementia as a route to such counterfactual comparisons^6,7^.

For research on cognitive reserve, a digital twin brain has to meet four requirements. The first, multitask dynamics, is to generate whole-brain signals during multiple cognitive tasks as well as at rest, because cognitive reserve concerns cognition. The second, individuality, is to reproduce these signals for an individual unseen during training, together with their individual-specific characteristics, so that the model generalizes to new people as eventual clinical use would require. The third, counterfactuals, is to alter conditions such as age while preserving that individual, reproducing faithfully the complex changes that these conditions bring about in the brain. The fourth, regional propagation, is to alter a single region and follow how the effect propagates to other regions through their connections, as the question of how regional contributions are reorganized demands.

Efforts to develop digital twin brains have followed two main lines of research, which meet complementary subsets of these requirements. Mechanistic models, most commonly neural mass models, assign each brain region a state that evolves according to biophysically grounded equations and couple these states through structural connectivity^8^, whereas data-driven models learn the patterns of brain signals directly from large-scale data. Multitask dynamics have been reproduced by data-driven foundation models^9^ but rarely by mechanistic models, whose generation has centered on resting-state activity^8^. Individuality is met only in part by both: mechanistic models, fitted to each individual by data assimilation of a few static parameters, reproduce individual-specific characteristics only to a limited extent^7,10–12^, and data-driven models reproduce individuals unseen during training only partly^13–15^.

Counterfactuals are likewise limited: mechanistic models implement aging with simple, uniform rules, such as removal of a fixed proportion of the connections between the hemispheres^8^, that may depart from the complex changes of the aging brain^16,17^, and data-driven models derive them only as group averages, without preserving the individual^18^. Regional propagation, finally, follows directly from the architecture of mechanistic models and has advanced to clinical trials in epilepsy^19^. Most data-driven models, by contrast, lack dynamics that unfold from region to region and from past to future: they generate each region’s signal directly from the observed signals of all regions and time points at once^20^ and are therefore not well suited to simulating how a change in one region spreads over time^21^. Although hybrid frameworks have begun to bridge the two lines^22–26^, the four requirements have so far been addressed largely in separate models.

Here we address the four requirements with Brain-VRNN, a Variational Bayesian Recurrent Neural Network (V-RNN) structured after the brain^27–29^, which generates brain signals from latent states evolving over time. For multitask dynamics, the model is trained on signals from multiple cognitive tasks and rest, and its latent states form a hierarchy from fast, region-specific to slow, whole-brain changes^29–32^, capturing both brief responses to task events and slower dynamics that persist throughout a task. For individuality, the roles are divided between training and test: during training on many individuals, the model learns the network weights that govern the dynamics shared across people, and at test, with these weights fixed, it estimates what is specific to an individual unseen during training by data assimilation of latent states that evolve over time.

Furthermore, for cognitive reserve, individuality has to be extended beyond a single observation: because the question of why the same aging leads to different degrees of decline is examined in longitudinal data, individuality comprises a trait-like part, stable within a person across time points, and a state-like part that varies between them. To capture individuality in longitudinal data, Brain-VRNN therefore disentangles these two parts into a trait dimension of its latent states, constrained during training to take a single value within a person across time points, and a state dimension, left free to vary between them.

For counterfactuals, conditions such as age and modifiable risk factors enter the model as inputs whose effects on the signals are learned from many individuals rather than postulated. Because individuality resides in the estimated latent states, shifting a condition while keeping these states fixed yields the signals of the same individual under that condition. For regional propagation, each of 52 regions holds its own state that evolves over time and is coupled to the others through structural connectivity, so that aging a single region alone, which is impossible in the real brain where all regions age together, propagates to the others and isolates that region’s contribution to the aging-related decline.

The Results follow the same order. We first show multitask dynamics and individuality together: the estimated latent states regenerate the multitask and resting-state signals of individuals unseen during training while preserving their individuality, both in the signals themselves and in four functional aging markers computed from them, which quantify brain functions that decline with age. For counterfactuals, counterfactual aging reproduced the aging-related decline of these functions while preserving individuality. For regional propagation, focal aging repeated across all 52 regions showed that the regions driving this decline differ across the markers, which addresses how regional contributions are reorganized with age. Finally, combining the four requirements, we return to the question of why the same aging leads to different degrees of decline, and show that the trait dimension, more stable across time points than the state dimension, contributed to the prediction of future cognitive change and modulated both the decline under counterfactual aging and the recovery obtained by an intervention on a modifiable risk factor^2^.

## RESULTS

### System overview

We applied Brain-VRNN to the Dallas Lifespan Brain Study (DLBS)^33^, a longitudinal cohort of 295 healthy adults followed for up to three time points (Methods). At each time point, fMRI was recorded during four tasks, memory encoding of visual scenes, semantic judgment at two levels of difficulty, passive viewing of visual categories, and rest, and the blood-oxygen-level-dependent (BOLD) signals were parcellated into 140 regions of interest (ROIs). Each person also provided a structural connectivity matrix derived from their own diffusion MRI, and condition variables: age, body mass index (BMI), blood pressure, and a regional map of amyloid burden from positron emission tomography (PET).

Brain-VRNN generates the BOLD from three levels of latent states (Fig. 1a). At the bottom, the 140 ROIs are grouped into 52 brain regions, each assigned a module with its own state that evolves over time; the modules are connected with weights fixed to the person’s structural connectivity, so that a change in one region propagates to the others, and each module generates the BOLD of the ROIs it covers. Above them, a single contextual module with a slower time constant sends a common input to all 52 regions, carrying whole-brain dynamics. At the top, a two-dimensional global latent summarizes a whole scanning run and therefore does not change within the run; its two dimensions are the trait and state dimensions into which individuality is disentangled. The condition variables and the time course of the task stimuli enter the contextual module and every regional module, so that a condition can be shifted either throughout the brain or in a single region.

**Fig 1.**
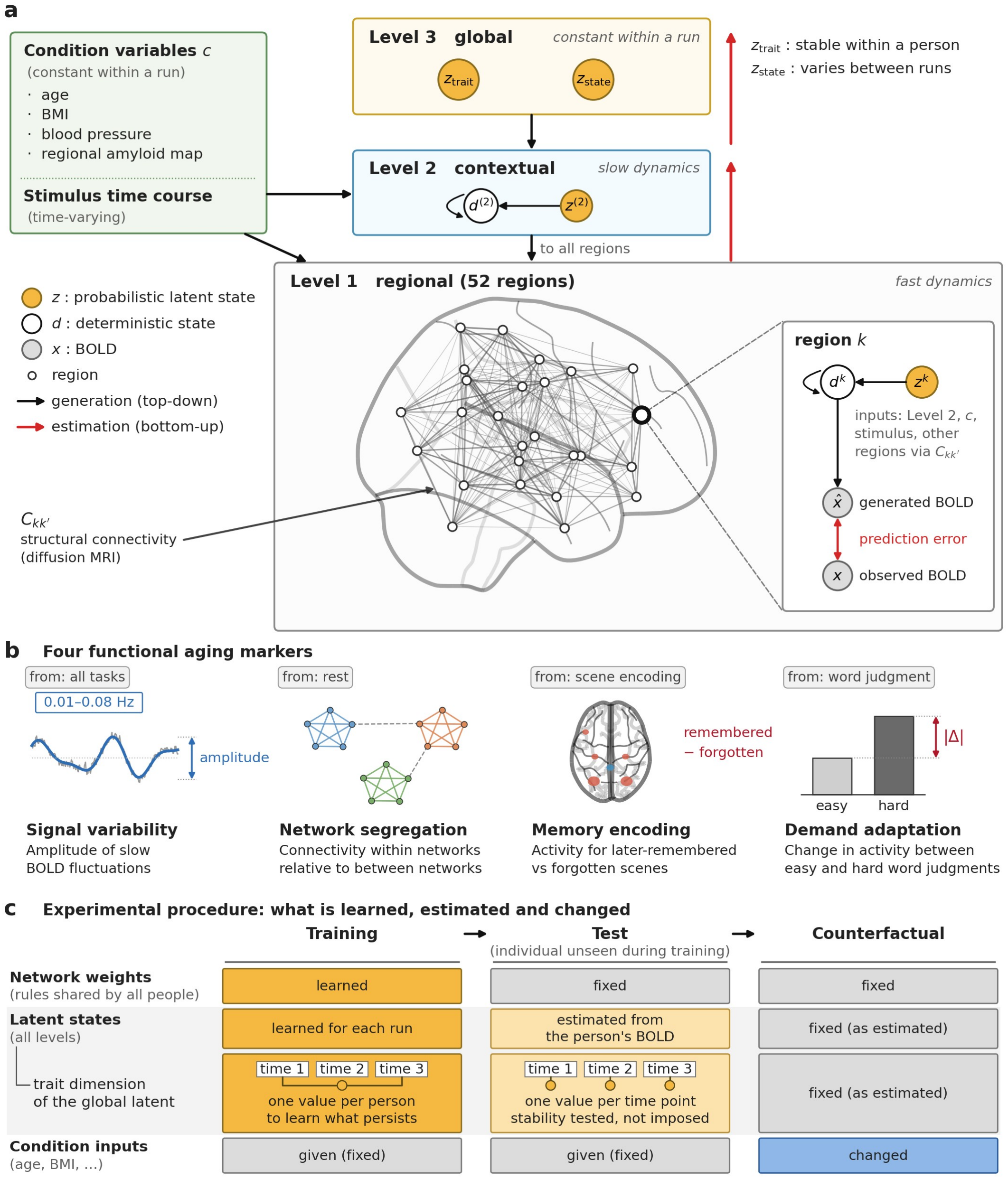
Brain-VRNN, the functional aging markers, and the experimental procedure. **a**, Brain-VRNN, a brain-structured hierarchical generative model. Three levels of latent states generate the BOLD of the four tasks from the top down: 52 regional modules (level 1), one per brain region (right hemisphere shown), coupled through lateral connections *C*_*k k ′*_ fixed to the person’s structural connectivity from diffusion MRI (lines, illustrative); a single contextual module (level 2) with slower dynamics that sends top-down input to all 52 regions; and a two-dimensional global latent (level 3), constant within a run, whose dimensions are the trait and state dimensions. The condition variables and the stimulus time course enter the contextual module and all regional modules. Inset, one regional module: its deterministic state *d*^*k*^ and latent state *z*^*k*^ generate the BOLD of the ROIs it covers, and prediction errors between the generated and observed BOLD are propagated bottom-up to estimate the latent states (red arrows). Unit numbers and time constants are given in Methods. **b**, The four functional aging markers: signal variability, the amplitude of slow (0.01–0.08 Hz) BOLD fluctuations; network segregation, connectivity within relative to between functional networks at rest; memory encoding, activity for later-remembered versus forgotten scenes; and demand adaptation, the change in activity between easy and hard word judgments in either direction (Methods). **c**, Experimental procedure: what is learned, estimated, kept fixed, and changed in training, at test on individuals unseen during training, and in the counterfactual experiments. The trait dimension takes one value per person across all time points in training, so that the model learns what persists within a person, and one value per time point at test, so that its stability across time points is tested rather than imposed. BMI, body mass index; ROI, region of interest.

To evaluate the BOLD in terms of brain function, we summarized it with four functional aging markers, each quantifying an aspect of brain function that declines with age, and each computed in the same way from the observed and the generated BOLD (Fig. 1b; Methods). Signal variability is the amplitude of the slow fluctuations of regional signals across all tasks; network segregation is how distinctly the brain divides into functional networks at rest; memory encoding is how strongly the brain expresses the activity pattern that predicts later remembering; and demand adaptation is how much brain activity changes when a task becomes harder.

The experimental procedure consists of three steps (Fig. 1c): training, test, and counterfactual experiments. In training, the model learns from the training individuals the network weights that govern the brain dynamics common to all people, together with the latent states of each of their runs. To implement the distinction between the trait-like and state-like parts, the trait dimension is constrained to take a single value for each person, the same in all of that person’s runs and time points, whereas the state dimension takes its own value in each run; the model thereby learns to read from the BOLD a component that persists within a person and a component that changes between time points. At test, the learned weights are fixed, and only the latent states of an individual unseen during training are estimated from that person’s BOLD, given their structural connectivity and condition values. If this estimation captures the person, the BOLD generated from these latent states should reproduce that person’s signals and functional aging markers in all tasks, including rest, and resemble them more closely than those of other people, preserving the person’s individuality. Unlike in training, the trait dimension is estimated separately at each time point; if it captures what persists within a person, its values should still agree across that person’s time points. In the counterfactual experiments, only the condition inputs are changed while the estimated latent states are kept fixed, and the markers of the regenerated BOLD are compared with those regenerated without that change, so that its effect is measured within the same person; if the model has learned how age acts on the brain, aging should shift the markers in the direction of the observed aging-related decline. Training and test individuals never overlap (individual-level 2-fold cross-validation; Methods).

### Brain-VRNN reproduces the multitask BOLD and individuality of unseen individuals

From the latent states estimated for individuals unseen during training, Brain-VRNN regenerated their BOLD in all tasks, including rest, and this BOLD met the first two requirements, multitask dynamics and individuality. The generated BOLD closely followed the observed time course (median r = 0.69, IQR 0.60 to 0.76; Fig. 2a), reproducing task-related fluctuations such as the fall of the signal during fixation periods. Behind this reproduction, the levels of the latent hierarchy played distinct roles: the regional states changed over time in a manner specific to each region, the contextual state supplied an input shared by all 52 regions, and the global latent remained constant within the run. The distribution of task responses across the 140 ROIs was also reproduced: the group-mean maps of the contrast between later-remembered and forgotten scenes and of that between hard and easy word judgments, estimated in the same way from the generated and the observed BOLD (Methods), correlated at r = 0.92 and r = 0.97 (Fig. 2b).

**Fig 2.**
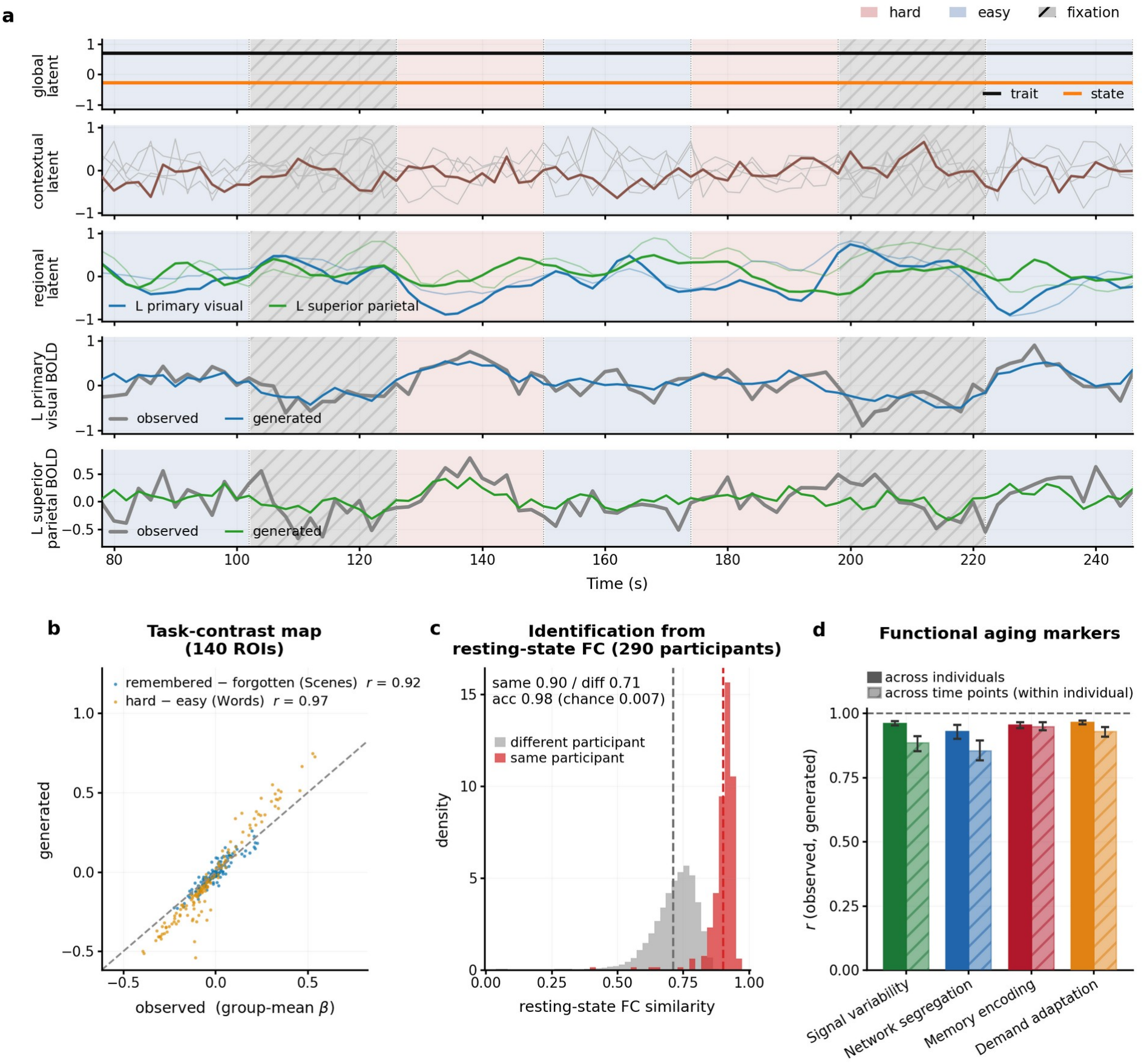
Brain-VRNN reproduces the multitask BOLD and individuality of unseen individuals. **a**, Representative example, a segment of one run of the semantic judgment task from a test individual: the posterior means (μ in Eq. 5) of the latent states at the three levels (top three rows; at the global level the trait and state dimensions, constant within the run; at the contextual level one of the five dimensions is highlighted; at the regional level two of the 52 regions are shown with their second dimension as thin lines) and the observed and generated BOLD of one region of interest (ROI) in each of the same two regions (bottom two rows); L, left. **b**, Group-mean task-contrast maps from the generated BOLD against those from the observed BOLD. For every test run, the same general linear model with the canonical hemodynamic response function was fitted to the generated and to the observed BOLD, giving the contrast β (remembered − forgotten for Scenes, hard − easy for Words) in each of the 140 ROIs, which was then averaged over runs; each point is one ROI; values, Pearson r across ROIs; dashed line, y = x. **c**, Similarity (Pearson correlation of the FC values) of the resting-state functional connectivity (FC) computed from the generated BOLD to the observed FC of the same participant or of other participants, pooled over the two folds (290 test participants); dashed lines, means; acc, identification accuracy averaged over the folds, with the chance level. **d**, Correlation between the four functional aging markers (Fig. 1b) computed from the observed and the generated BOLD, across individuals (solid bars) and across time points within an individual (hatched bars); error bars, 95% CI from a bootstrap over participants.

The generated BOLD also preserved each person’s individuality, both at rest and during the tasks. At rest, functional connectivity (FC) computed from the generated BOLD correlated with the person’s own observed FC at r = 0.90, exceeding the correlation with other people (r = 0.71), so that each person was identified among the test participants of the same fold with an accuracy of 0.98 (chance = 0.007; Fig. 2c). During the tasks, individuality extended to the four functional aging markers (signal variability, network segregation, memory encoding, and demand adaptation; Fig. 1b), three of which are computed from task BOLD: the markers computed from the generated BOLD correlated with the observed values at r = 0.93 to 0.96 across individuals and r = 0.86 to 0.95 across time points within the same individual (Fig. 2d). These individual differences also carried the same cognitive meaning: whether computed from the generated or the observed BOLD, the markers showed nearly the same pattern of correlations with the scores of 16 cognitive tests obtained outside the scanner (r = 0.99 between the two patterns; Supplementary Tables 1 to 4).

### The global latent separates into trait and state dimensions

Within this individuality, the two dimensions of the global latent separated what persists within a person from what changes between time points. Although the trait dimension was estimated separately at each time point, its values agreed across time points within the same individual (r = 0.58 [0.45 to 0.67]; Fig. 3a), substantially more than those of the state dimension (r = 0.14 [0.08 to 0.20]; Fig. 3b; Δr = +0.44 [+0.32 to +0.54], p < 0.0001). The trait dimension likewise mainly expressed between-individual variance whereas the state dimension mainly expressed within-individual variance (intraclass correlation coefficient, ICC = 0.78 [0.72 to 0.83] versus 0.29 [0.22 to 0.38]; ΔICC = +0.48 [+0.39 to +0.57], p < 0.001). The model thus acquired in training the ability to separate, from the BOLD of a single time point, what will persist in a person from what will change.

**Fig 3.**
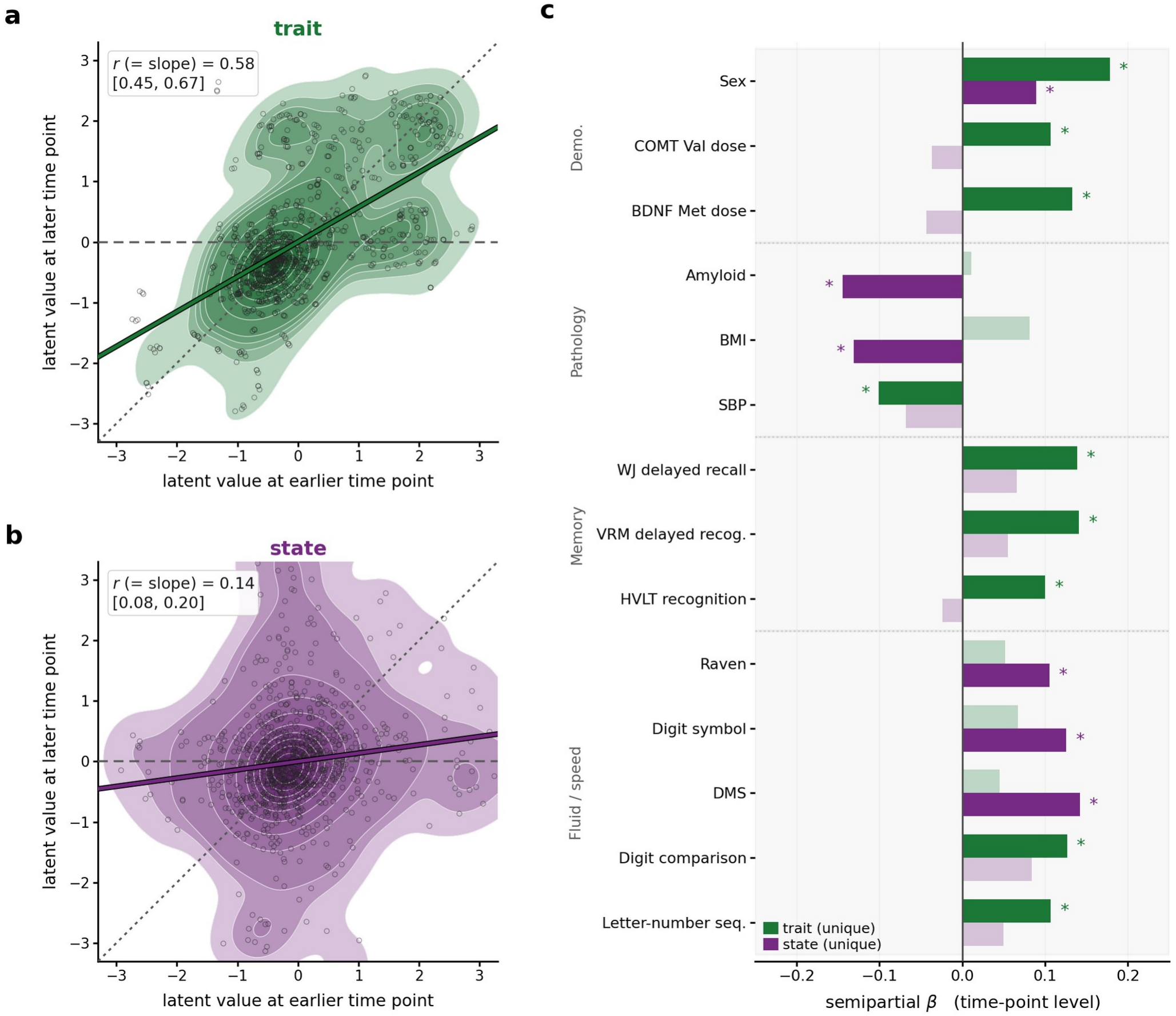
The global latent separates into trait and state dimensions. **a**, Test-retest of the trait value between an earlier and a later time point of the same test individual. Each point is one individual, one pair of time points, and one task; values are z-scored across individuals within each task and pair of time points. Shading, density layers covering 10 to 99% of the points; solid line, regression slope; inset, r with its 95% CI from a bootstrap over participants; dotted line, y = x; dashed line, y = 0. **b**, The same for the state value. **c**, Unique association of the trait (green) and state (purple) values with external variables, expressed as the semipartial standardized β of each value after adjusting for the other. Only variables with p < 0.05 for at least one of the two values are shown. Saturated bars with asterisks, p < 0.05; pale bars, not significant. Demo., demographic and genetic variables; COMT Val dose, number of valine alleles of the catechol-O-methyltransferase gene; BDNF Met dose, number of methionine alleles of the brain-derived neurotrophic factor gene; BMI, body mass index; SBP, systolic blood pressure; WJ, Woodcock–Johnson; VRM, verbal recognition memory; recog., recognition; HVLT, Hopkins verbal learning test; DMS, delayed matching to sample; seq., sequencing; VRM and DMS are from the Cambridge Neuropsychological Test Automated Battery.

The two dimensions were also associated with different external variables (Fig. 3c). As expected, measures that are invariant within a person, such as sex and genotype, were associated with the trait value, whereas measures that vary within a person, such as amyloid burden and BMI, were associated with the state value. Among the cognitive test scores, all three episodic memory measures were associated with the trait value alone, whereas the measures associated with the state value alone came from reasoning, processing speed, and working memory.

### Counterfactual aging reproduces the aging-related decline while preserving individuality

Brain-VRNN also met the third requirement, counterfactuals: when only the age condition was shifted, the four functional aging markers computed from the generated BOLD moved in the direction of the observed aging-related decline (Fig. 4). To define this reference direction, we first measured the aging-related decline itself: across the test individuals and their time points, all four markers were lower at older ages in the observed BOLD, and the BOLD generated from the estimated latent states at each person’s actual age reproduced this association with age at 86 to 99% of its observed size (Fig. 4a). We then kept these latent states fixed, shifted only the age condition within each person, and generated the BOLD again; every marker computed from this BOLD decreased, consistent with the observed aging-related decline (Fig. 4b; its size is considered in the Discussion).

**Fig 4.**
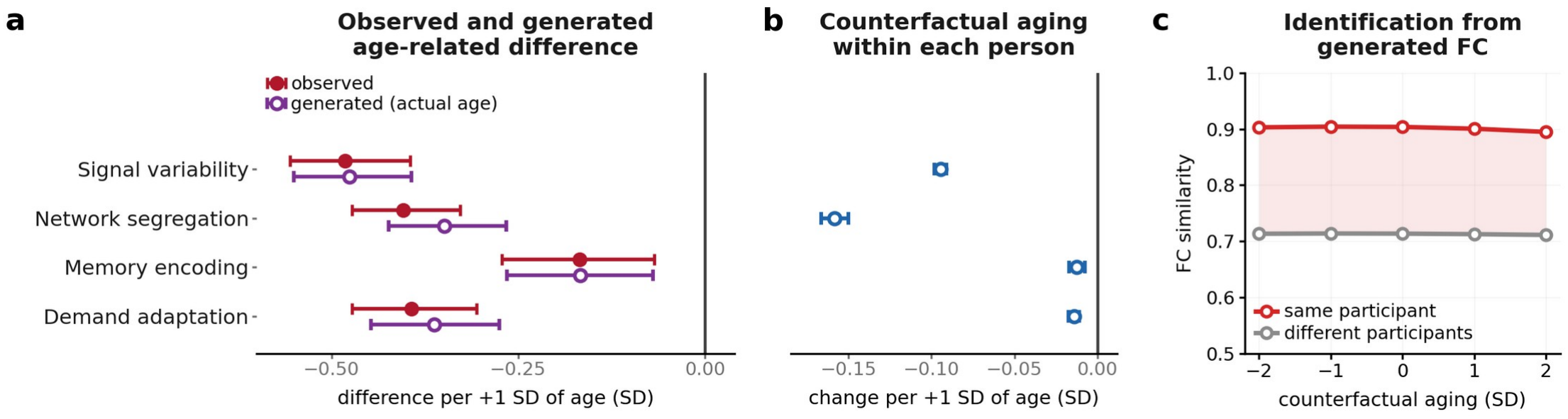
Counterfactual aging reproduces the aging-related decline while preserving individuality. **a**, Age-related difference of the four functional aging markers computed from the observed BOLD (filled) and from the BOLD generated from the estimated latent states at each person’s actual age (open), expressed as the regression slope on age across all sessions of the test individuals, pooling individuals and time points (SD of the marker per +1 SD of age). **b**, Change in the markers induced by counterfactual aging, in which only the age condition is shifted by +1 SD with the estimated latent states kept fixed; note the different scale from **a**. In **a** and **b**, circles and bars denote the estimate and its 95% CI from a bootstrap over participants, and negative values indicate a decrease with age. **c**, Similarity of resting-state functional connectivity (FC) computed from the BOLD generated under counterfactual aging from −2 to +2 SD to the observed FC of the same participant (red) and of different participants (gray), computed as in Fig. 2c.

The counterfactual aging also preserved individuality (Fig. 4c). Across doses from −2 to +2 SD of age, the FC computed from the generated BOLD stayed correlated with the person’s own observed FC at about 0.90, above the correlation with other people (about 0.71), and individual identification remained at an accuracy of 0.98.

### The regions that drive the aging-related decline differ across functional aging markers

The fourth requirement, regional propagation, was then used to ask how much each brain region contributes to the aging-related decline of the four markers. When a single region is counterfactually aged, the effect propagates from that region to the others through their connections, so that it can be evaluated as the change in the markers computed from the BOLD of the whole brain. Repeating this over the 52 regions therefore isolates the contribution of each region to the decline. In this virtual focal aging (Fig. 5a), one region was aged while the other 51 remained at the person’s actual age, and the contribution was given by the change in each marker relative to the generation with no region aged. Focal aging was complemented by virtual focal sparing, in which every region was aged except one, and the change relative to the generation with every region aged gave how much sparing that region alone recovers the markers. The two simulations thus differ both in the age of the other regions and in the direction of the change, and the agreement between the regions they identified was examined.

**Fig 5.**
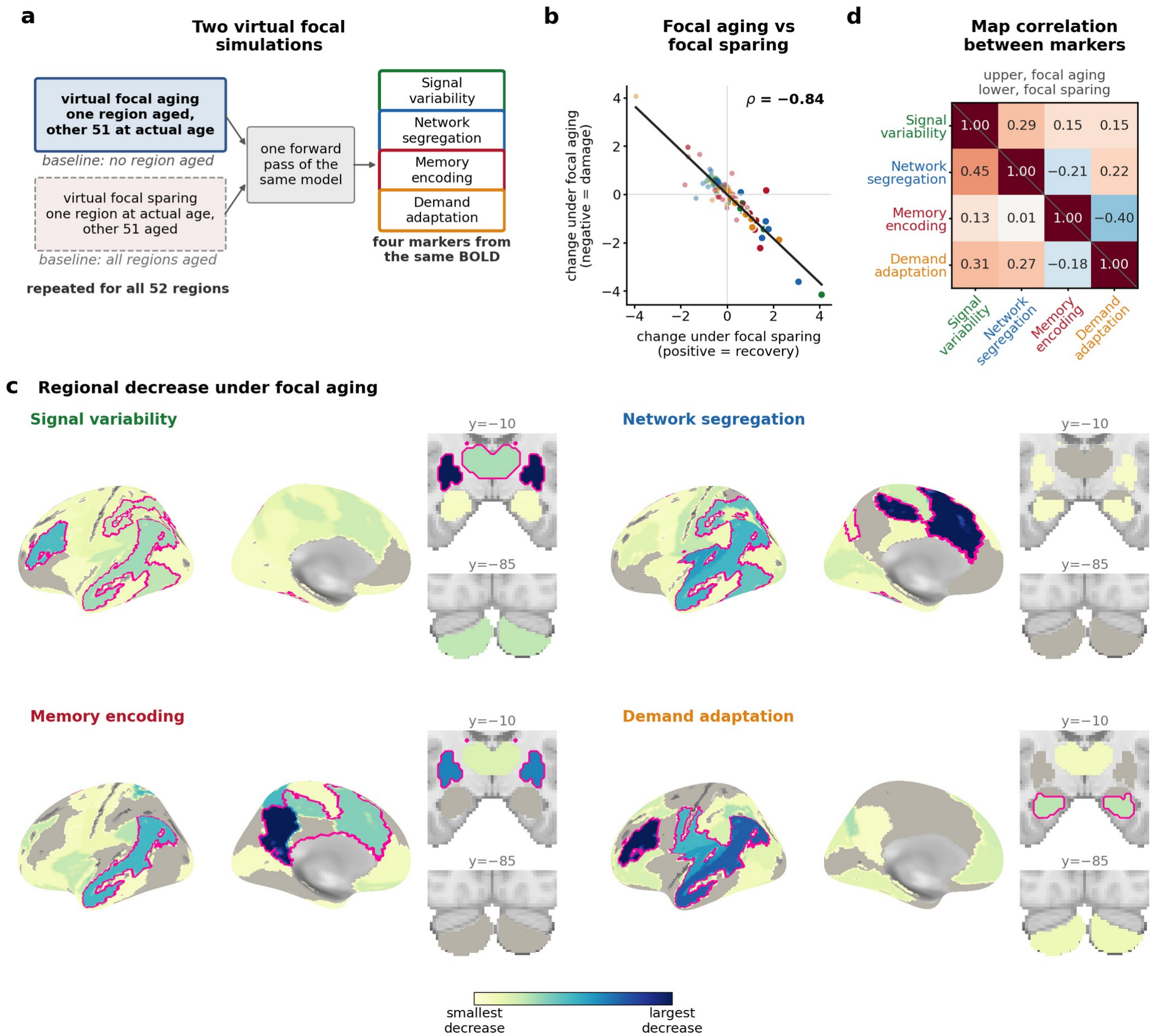
The regions that drive the aging-related decline differ across functional aging markers. **a**, The two virtual focal simulations. In focal aging, one region is aged and the other 51 remain at the person’s actual age; baseline, no region aged. In focal sparing, one region remains at the actual age and the other 51 are aged; baseline, every region aged. Both are repeated over the 52 regions, and the four markers are computed from the same generated BOLD. Colors denote the four markers throughout the figure. **b**, Change under focal aging (*Δ*_*aging*_, negative for damage) against change under focal sparing (*Δ*_*sparing*_, positive for recovery) for the 26 regions of each marker (homologous regions of the two hemispheres combined, z-scored within each marker), colored by marker. Line, fit pooled over the four markers; ρ, Spearman correlation over the four markers combined (per marker: signal variability −0.86, network segregation −0.73, memory encoding −0.76, demand adaptation −0.76). **c**, Regional maps of *Δ*_*aging*_, the decrease of a marker when that region alone is aged. Colors are scaled within each marker from the smallest (light) to the largest (dark) decrease, so the color scale is not comparable between markers; gray, regions whose marker did not decrease. Homologous regions share one value; the left hemisphere is shown (lateral and medial surfaces) with coronal sections at y = −10 and y = −85 for the subcortex and cerebellum. Magenta outlines, the five regions (top 20%) with the largest *Δ*_*s par in g*_. **d**, Spearman correlation between the regional maps of the four markers under focal aging (upper triangle) and under focal sparing (lower triangle), computed on the centered effects.

The two simulations identified largely the same regions. Their regional maps were strongly negatively correlated (ρ = −0.84 across the four markers combined; Fig. 5b): the regions whose aging lowered a marker were those whose sparing recovered it. This correspondence held for single regions as well: across the four markers, 85% of the five regions with the largest decrease under focal aging were also among the five with the largest recovery under focal sparing (Fig. 5c, magenta outlines).

In contrast, the regions that drove the decline differed from one marker to another: the regional maps did not reach high positive correlations between markers in either experiment (at most +0.29 for focal aging and +0.45 for focal sparing; Fig. 5d). These low correlations do not reflect noise: each map was reproduced almost exactly between two non-overlapping halves of the individuals (r = 0.94 to 1.00). In the focal-aging maps (Fig. 5c), signal variability depended most strongly on the striatum/globus pallidus, the dorsolateral prefrontal cortex, and the thalamus, corresponding to a fronto-striato-thalamic system. Network segregation depended most strongly on the dorsal anterior cingulate cortex and pre-supplementary motor area, and on the mid-posterior cingulate, the medial wall of the salience/ventral attention system. Memory encoding depended most strongly on the posterior cingulate cortex and precuneus, and on the striatum/globus pallidus. Demand adaptation depended most strongly on the dorsolateral prefrontal cortex and on a region spanning the lateral temporal cortex and angular gyrus. Although the four markers depended on largely distinct regions, one region ranked among the top five for every marker in both simulations: the region spanning the lateral temporal cortex and the angular gyrus, a candidate aging hub shared across the markers.

### The trait dimension predicts individual differences in aging-related decline

With the four requirements met, we returned to the question posed at the outset, why the same aging leads to different degrees of decline in different individuals. To address this question, we focused on the trait dimension of the global latent, which expresses the part of individuality that remained stable within a person across time points (Fig. 3a). We hypothesized that the trait value is related to how cognition and brain function change with aging. To test this hypothesis, we first examined in the observed longitudinal data whether the trait value estimated at one time point predicted the scores of the cognitive tests at the next time point beyond their current values (Methods). A higher trait value predicted higher future scores of the Mini-Mental State Examination and of tests of episodic memory and reasoning, three of the 12 tests, although the effects were small (standardized β +0.07 to +0.15, q = 0.026; Fig. 6a). As expected, the state value predicted none of them, indicating that the two dimensions represent separate components of individuality.

**Fig 6.**
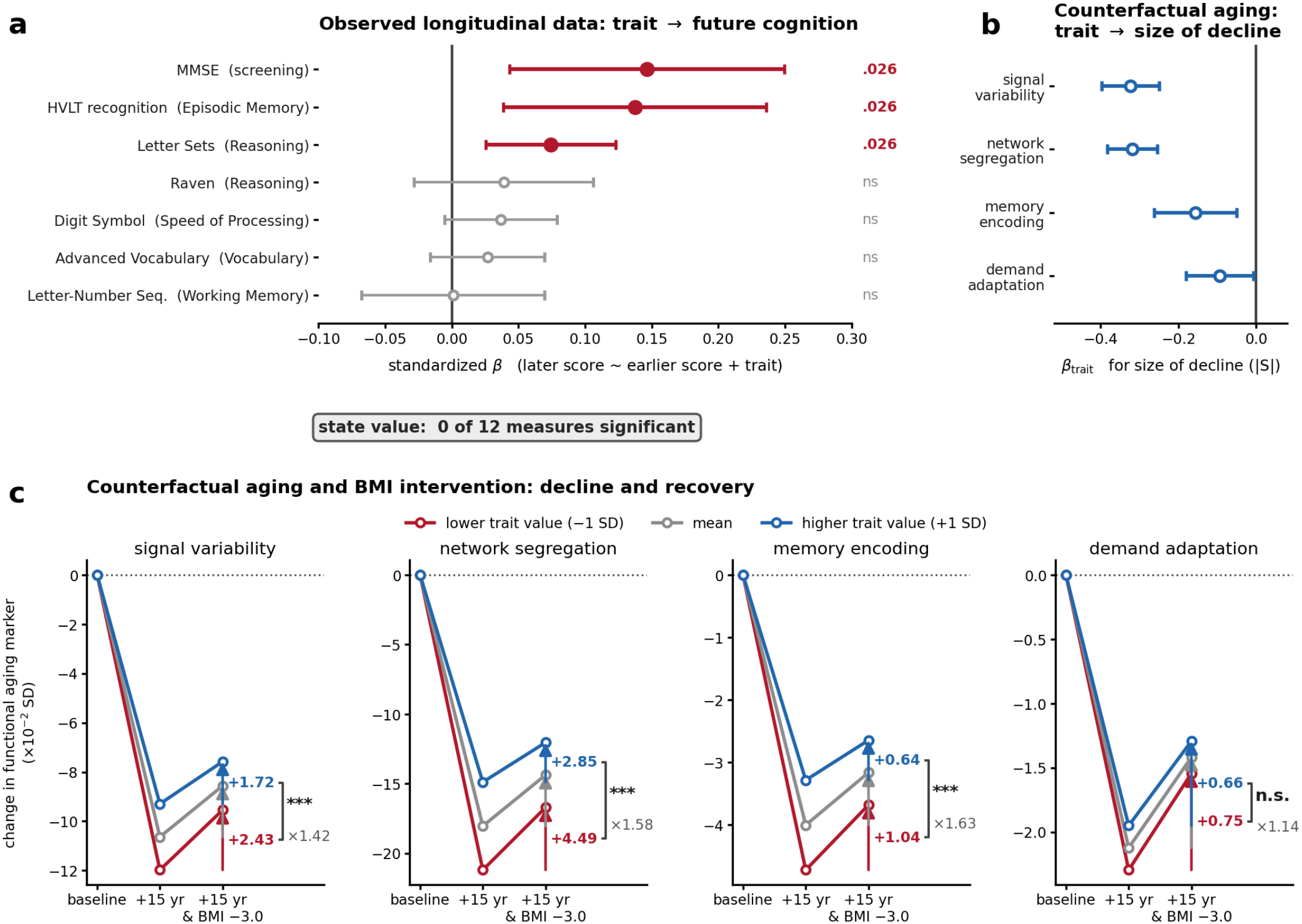
The trait dimension predicts individual differences in aging-related decline and in the recovery obtained by a simulated intervention. **a**, Prediction of future cognition by the trait value in the observed longitudinal data: standardized β of the trait value in the model of the later score on the earlier score and the trait value, with the trait value estimated from the BOLD of the earlier time point only. Seven of the 12 measures are shown; filled circles, q < 0.05 after Benjamini–Hochberg correction over the 12 measures; error bars, 95% CI. MMSE, Mini-Mental State Examination; HVLT, Hopkins verbal learning test; Seq., sequencing. **b**, Association between the trait value and the size of the decline of each functional aging marker under counterfactual aging (β of the trait value for |S|, Eq. 10); negative values indicate a smaller decline at a higher trait value; the sign of the trait dimension was fixed by a convention independent of these outcomes (Methods). Error bars, 95% CI. **c**, Change in each functional aging marker (×10^−2^ SD) relative to the baseline generation at each individual’s actual age, under aging alone (+15 years, +1 SD) and under the same aging combined with a reduction of body mass index (BMI) by 3.0 kg/m^2^, for individuals with a lower (1 SD below the mean; red), average (gray), and higher (1 SD above the mean; blue) trait value, predicted from the regression on the trait value (Methods). Numbers, recovery R (Eq. 12); ×N, ratio of the recovery for a lower to that for a higher trait value; ***, p < 0.001; n.s., not significant.

Next, we examined in the model whether the trait value was likewise related to the decline of brain function under counterfactual aging, in which every individual was aged by the same amount (Methods). Under counterfactual aging, individuals with a higher trait value showed a smaller decline in all four functional aging markers (*β*_*trait*_ = −0.32 for signal variability, −0.32 for network segregation, −0.16 for memory encoding, and −0.09 for demand adaptation), with none of the 95% CIs crossing zero (Fig. 6b). The trait value was thus related to how both cognition and brain function change with aging, in the observed course and under counterfactual aging alike. Such individual differences in how much a person is affected by aging are what the concepts of cognitive reserve and, more broadly, resilience to aging were proposed to explain^2^.

### The trait dimension also predicts individual differences in the recovery obtained by a simulated intervention

For prevention, it matters not only who declines more with aging but also who would benefit from an intervention. We therefore asked whether the trait value, which marked the individuals who decline more (Fig. 6b), also relates to how much of this decline an intervention can reverse. To answer this, we first simulated an intervention and measured its average effect, and then examined whether this effect depended on the trait value. Of the modifiable risk factors available in this general-population cohort (Methods), only BMI was associated with the functional aging markers in a consistent direction after age adjustment in the observed data, albeit weakly, and BMI was therefore chosen as the target of the intervention (for the other conditions, see Supplementary Fig. 1 and the Discussion). The intervention was then simulated in the model in the same way as the counterfactual aging, with each individual’s latent states kept fixed and only the condition inputs changed (Fig. 6c; Methods). A reduction of BMI by 3.0 kg/m^2^ (0.91 SD) was combined with aging by +15 years (1 SD), and the BOLD regenerated under this combination was compared with that regenerated under the same aging alone. The difference between the two isolates the effect of the intervention within the same individual under the same aging, a comparison that observational data cannot provide. On average, the reduction of BMI reversed about one fifth of the decline caused by the aging (20 to 33% across the four markers; Fig. 6c, gray).

We then asked whether this recovery differed with the trait value. Consistent with Fig. 6b, individuals with a lower trait value declined more under the aging (Fig. 6c; red and blue, 1 SD below and above the mean trait value). Individuals with a lower trait value also recovered more with the reduction of BMI, their recovery being 1.42, 1.58, and 1.63 times that of individuals with a higher trait value for signal variability, network segregation, and memory encoding (all p < 0.001), although the difference did not reach significance for demand adaptation (1.14 times). This relation was not explained by a ceiling effect (Methods). The trait value thus captured not only each individual’s vulnerability to aging but also the recovery obtained by an intervention, which was larger in the individuals with a lower trait value.

## DISCUSSION

This study asked whether a digital twin brain could meet the four requirements that research on cognitive reserve places on a model, and used it to ask why the same aging leads to different degrees of decline. Brain-VRNN regenerated the multitask BOLD of individuals unseen during training while preserving their individuality (Fig. 2), and separated this individuality into a trait dimension stable within a person and a state dimension varying across time points (Fig. 3). Because the effects of the conditions are learned from many individuals and individuality resides in the estimated latent states, counterfactual aging reproduced the aging-related decline while preserving the individual (Fig. 4). Because each region holds its own state coupled through structural connectivity, focal aging repeated over the 52 regions showed that the regions driving the decline differ across the functional aging markers (Fig. 5). Combining the four requirements, the trait value predicted the future course of cognition, the decline under counterfactual aging, and the recovery obtained by a simulated intervention (Fig. 6).

Which regions drive the decline of whole-brain function with age is difficult to establish from correlational analyses of observational data. The focal aging and sparing experiments address this question directly. In these experiments, the regions that drive the decline differed across the four functional aging markers rather than following a single deterioration, in line with the many distinct modes into which brain aging decomposes in large-scale data^34^ and with the non-uniformity of dedifferentiation across regions and tasks^5,35^. Moreover, the regions that drove the decline of each marker match where the corresponding change with age has been localized: the fronto-striato-thalamic system for signal variability^36,37^, the posterior cingulate cortex and precuneus for memory encoding^38,39^, the dorsolateral prefrontal cortex for demand adaptation^4,40,41^, and the medial wall and association cortices for network segregation^42^. Across this separation, a single region spanning the lateral temporal cortex and the angular gyrus ranked high for all four markers, consistent with accounts of connectivity hubs as sites of vulnerability and of the angular gyrus as an integrative hub^43–45^.

Notably, no cognitive score was used in training: the trait dimension was constrained only to take the same value across a person’s time points, not to predict cognition or decline. Nonetheless, this dimension, which the model needed only to generate an individual’s BOLD, predicted both the future course of cognition and the decline under counterfactual aging (Fig. 6a,b), the kind of individual difference that cognitive reserve was proposed to explain^2^. This dimension shares some elements with the concept of cognitive reserve, being a characteristic of the person estimated from functional brain activity common to multiple tasks, as proposed for neural measures of reserve^2^. It differs in others: it was not positively associated with education, a common proxy of reserve (r = −0.10), and whether it moderates the effect of brain changes on cognition, as reserve is defined to do, was not tested. Because it also related to a smaller decline of brain function itself, it may be viewed in terms of the broader notion of resilience to aging, which includes brain maintenance as well as reserve^2,3^.

Several limitations follow from the data and from the model. DLBS is a sample of healthy adults across the adult lifespan, so that the age axis mainly reflects healthy aging, and the pathological processes that lead to cognitive impairment are beyond what the present data can address. Blood pressure and amyloid burden, both supplied to the model as conditions, did not produce a consistent recovery when reduced, unlike BMI (Supplementary Fig. 1); since neither was associated with the functional aging markers in the observed data, this may likewise reflect a sample close to health, in which the interaction between reserve and pathology is difficult to detect^46,47^. Within-person change is also sparsely sampled, with about 4 years between time points and three time points for part of the cohort. On the model side, the size of the change that counterfactual aging can produce may be limited, because only the conditions are shifted while the latent states and the learned weights stay fixed. The claims made here therefore concern the direction of change and the differences between individuals rather than absolute magnitudes. The responses to intervention are likewise generated: the shifts were kept within a realistic range for this cohort (Methods) and the response to BMI agreed in direction with its observed association, but the response of an individual to an actual intervention could not be validated, because DLBS includes no intervention.

The limitations that follow from the data would be relaxed by training on a cohort that includes cognitive decline of pathological origin. In such a cohort, the interaction between reserve and pathology becomes testable. The architecture places no restriction on which conditions can be shifted as interventions, since any measured condition enters by the same route as BMI does here, so that risk factors for dementia beyond BMI can be expected to take effect in such data. The response of one individual to each intervention can then be generated and compared, and the resulting ranking is what an individualized choice of intervention would rest on^48,49^. A generative brain model trained on such data could thus advise on the prevention of cognitive decline for a person from interventions simulated for that same person.

## METHODS

### Participants and data

This study used the Dallas Lifespan Brain Study (DLBS), a publicly available longitudinal dataset of healthy, community-dwelling adults followed for up to three time points at intervals of roughly 4 years (3–5 years)^33^. Throughout, a session denotes all scans of a person at one time point, and a run denotes one continuous acquisition of one task within a session; runs are the unit that the model generates. Sessions were included when task and resting-state fMRI, diffusion MRI, and PET (amyloid or tau) were available, together with APOE genotyping, which yielded 295 participants (181 women) and 535 sessions (262, 151, and 122 sessions at time points 1 to 3); 131 participants contributed one session, 88 two, and 76 three. Age at scanning ranged from 30 to 96 years (median 68, IQR 58 to 76).

Each session comprised up to four fMRI tasks, each acquired at one volume every 2 s (repetition time): memory encoding of visual scenes (Scenes; three runs of 171 volumes), semantic judgment of words at two difficulty levels (Words; one run of 231 volumes), passive viewing of visual categories (VentralVisual; two runs of 202 volumes, acquired at time points 1 and 2 only), and rest (two runs of 154 to 210 volumes). In Scenes, participants viewed 96 outdoor scenes (32 per run) and judged whether water was present, an incidental encoding task; each scene was shown for 3 s with intertrial intervals jittered between 4 and 14 s, and each trial was classified by a recognition test given about 20 min later outside the scanner, with 96 lures, as remembered (with high or low confidence) or forgotten. In Words, participants judged whether each word denoted a living thing; 24-s blocks of eight words, eight blocks of semantically ambiguous words (hard; for example, “virus”) and eight of unambiguous words (easy; for example, “asphalt”), were presented in pseudorandom order with three 24-s fixation blocks as a baseline. VentralVisual presented 16-s blocks of faces (human and animal), houses, chairs, and scrambled images, and rest was acquired awake with a fixation cross; VentralVisual contributes to no functional aging marker other than signal variability, which is computed over all four tasks. Full descriptions of the tasks are given in the data descriptor^33^.

The participants were split into two halves at the individual level, so that all sessions of a participant fell on the same side, with the number of time points, sex, age, Mini-Mental State Examination (MMSE) score, and amyloid status balanced between the halves. The two halves served in turn as training and test sets (2-fold cross-validation): fold 1 trained on 148 participants (271 sessions, 1785 runs) and tested on 147 (264 sessions, 1692 runs), and fold 2 reversed the roles. Every result reported for test individuals therefore comes from a model that never saw those individuals during training, and the two folds together cover all 295 participants. The original study obtained written informed consent from all participants, and the present secondary analysis of deidentified data was approved by the Institutional Review Board of the National Center of Neurology and Psychiatry (IRB no. A2021-119).

### MRI acquisition and preprocessing

Functional images were acquired on a 3-T Philips scanner with a repetition time of 2 s and an echo time of 25 ms. The images were preprocessed with fMRIPrep (version 24.1.1)^50^ and normalized to MNI152NLin2009cAsym space. Regional time series were obtained by averaging voxels within 140 regions of interest (ROIs): 100 cortical parcels of the Schaefer atlas^51^ assigned to seven networks^52^, 32 subcortical parcels of the Tian atlas (scale II)^53^, and 8 cerebellar parcels of the Diedrichsen atlas^54^. The following confounds were regressed out of each ROI time series: the global signal, framewise displacement, six motion parameters and their derivatives, the first six anatomical CompCor components^55^, and three cosine regressors implementing a high-pass filter at about 1/128 Hz. No spatial smoothing or band-pass filtering was applied, and volumes were not censored so that each run remained a continuous time series for the model. For the model, the 140 ROIs were grouped into 52 brain regions defined as hemisphere by functional module: 22 cortical modules obtained by subdividing the seven networks (for example, dorsolateral prefrontal cortex, parietal cortex, and medial prefrontal cortex within the executive control network), plus the hippocampus, amygdala, thalamus, striatum and globus pallidus, and cerebellum, giving 26 modules per hemisphere with 1 to 8 ROIs each. Each ROI time series was linearly scaled within each run to the range of −0.9 to 0.9 to match the output range of the model.

### Structural connectivity

Diffusion-weighted images (30 directions at b = 1000 s/mm^2^ and one b = 0 image) were processed with MRtrix3^56^. The processing comprised denoising^57^, Gibbs-ringing removal^58^, eddy-current and motion correction (FSL eddy)^59^, bias-field correction, five-tissue-type segmentation of the T1-weighted image, multi-shell multi-tissue constrained spherical deconvolution^60,61^, and anatomically constrained probabilistic tractography^62^ with 10 million streamlines seeded at the gray matter–white matter interface (maximum length 250 mm, cutoff 0.06). Streamline weights were estimated with SIFT2^63^, and the connectome was calculated from the weighted streamlines connecting each pair of ROIs, symmetrized and aggregated to the 52 regions. The resulting matrix was log-transformed as log10(1 + x) and clipped at the mean ± 3 SD within each matrix; no consistency thresholding was applied, so that individual differences in weak connections were retained. A separate matrix was computed for every session, and the matrix of the corresponding session was supplied to the model as the structural connectivity C of that person, with the diagonal set to zero.

### Amyloid PET

Amyloid PET was acquired with 18F-florbetapir as a single 10-min frame starting 50 min after injection. The image was coregistered to the T1-weighted image and normalized to MNI space, and the standardized uptake value ratio (SUVR) was computed with the cerebellum (excluding the deep nuclei) as the reference region. Subcortical ROIs were eroded before sampling, and the striatum, globus pallidus, and thalamus were excluded because of nonspecific binding, leaving 48 of the 52 regions with a value of ln(SUVR). Two quantities were derived from these values, both standardized with training-set statistics only: a global amyloid burden and a regional amyloid deviation. The global amyloid burden was defined as the mean over ten regions of the Alzheimer signature (bilateral parietal, temporal, ventral and dorsal medial prefrontal, and posterior cingulate regions of the executive control and default mode networks) and transformed to a normal score by rank-based inverse-normal transformation. The regional amyloid deviation was defined for each region as ln(SUVR) minus the global burden, winsorized at the median ± 3 median absolute deviations and scaled by robust z-scoring. Runs without amyloid PET (7.9% of training runs and 9.9% of test runs) received zero for both quantities, the value of the training mean, so that these runs contributed no gradient to the amyloid pathway. Tau PET (18F-flortaucipir) was acquired in DLBS only in a subset of participants at time points 2 and 3, so that it covered 162 of the 535 sessions (30%, 139 participants) against 483 (90%) for amyloid; tau was therefore used only as an inclusion criterion and not supplied to the model.

### Condition variables and stimulus time course

Besides age, the axis of this study, the condition variables were BMI, systolic blood pressure, and amyloid burden. BMI and blood pressure represent midlife obesity and hypertension, two of the modifiable risk factors for dementia listed by the Lancet Commission^64^, which act on the brain through metabolic and vascular pathways, respectively; amyloid burden (Amyloid PET) is a pathological hallmark of Alzheimer’s disease. In the observed data, BMI was weakly associated with the functional aging markers in a consistent direction after age adjustment, whereas neither blood pressure nor amyloid burden was. Blood pressure and amyloid burden were nevertheless included because each represents a pathway to dementia distinct from the metabolic one represented by BMI: blood pressure the vascular pathway, and amyloid burden the amyloid pathology of Alzheimer’s disease. Age, BMI, and blood pressure were supplied to the model as constants within a session, age as the age at scanning and blood pressure as the mean of four readings on two days. Each was z-scored with the mean and SD of the training set (age 66.2 ± 14.8 years, BMI 25.8 ± 3.3 kg/m^2^, blood pressure 129 ± 18.0 mmHg). Amyloid burden was supplied as the two quantities derived in Amyloid PET, the global amyloid burden to the contextual module and the regional amyloid deviation to each brain region, so that the model receives how much amyloid a person carries and where it lies relative to that level.

The stimulus time course of the Words task was supplied as two binary channels indicating, at each volume, whether a hard or an easy block was presented; both channels were zero in the other tasks. This was the only stimulus input to the model. The model is generative rather than stimulus-driven, and task-evoked activity is reproduced from the latent states estimated for each run; a stimulus input was added only where an experimenter-controlled manipulation varies within a run and serves as an axis of counterfactual generation, which holds for the demand manipulation of Words alone. In Scenes, the quantity of interest is whether each scene is later remembered, an outcome of the participant rather than a manipulation, so that supplying it as an input would be circular; VentralVisual involves no parametric manipulation and rest none at all. The trial-locked activity of Scenes, and hence the subsequent memory effect, therefore has to be reproduced from the estimated latent states. The condition variables and the stimulus channels together form a six-dimensional condition vector for each module, *c*_*t*_ =(*age, BMI, BP, hard*_*t*_, *easy*_*t*_, *amy*), where the sixth element is the regional amyloid deviation for a brain region and the global amyloid burden for the contextual module.

### Cognitive test scores

Sixteen tests administered outside the scanner were used^33^: the MMSE; digit comparison and digit symbol (DigSym) for processing speed; spatial working memory errors, letter–number sequencing, and the CANTAB delayed matching to sample (DMS) and spatial recognition memory for working memory; Hopkins verbal learning test delayed recognition (HVLT), Woodcock–Johnson delayed recall (WJ), and the CANTAB verbal recognition memory (VRM) for episodic memory; Raven matrices, Letter Sets, and the CANTAB stockings of Cambridge for reasoning; the Educational Testing Service (ETS) vocabulary test and graded naming for vocabulary; and controlled oral word association for verbal fluency. Twelve of these were repeated at later time points and were used for the longitudinal analyses. A general cognitive score (PC1) was defined for Supplementary Tables 1 to 4 as the first principal component of the standardized scores of the MMSE and the nine tests with low missingness (that is, excluding the CANTAB DMS, spatial recognition memory, and VRM, controlled oral word association, graded naming, and Woodcock–Johnson delayed recall); sessions lacking five or more of these tests were excluded, the remaining missing entries (1.2%) were filled by iterative imputation with extra-trees regressors, and the component was fitted on the training half and projected onto the test half.

### Functional aging markers derived from BOLD

Four functional aging markers were computed from the observed BOLD and, by the same procedure, from the BOLD generated by the model described below; each was chosen to capture a functional change that earlier work has associated with aging and was defined so that it declines with age, two from the signal itself (signal variability and network segregation) and two from task-evoked responses (memory encoding and demand adaptation). Each ROI time series was z-scored within the run before computation, and session values were obtained by averaging over the runs of the session. The amplitude of BOLD fluctuation declines with age^36,37^; signal variability was therefore quantified by the fractional amplitude of low-frequency fluctuation (fALFF)^65^, computed for every run of the four tasks as the sum of the Fourier amplitudes between 0.01 and 0.08 Hz divided by the sum over the whole spectrum, averaged over the 140 ROIs. The segregation of functional systems decreases across the adult lifespan^42^; network segregation was therefore quantified by system segregation, computed from the resting-state runs following the original definition^42^: functional connectivity was computed as Fisher-transformed Pearson correlations between the 100 cortical ROIs with negative values set to zero, and system segregation was then (*Z*_*w*_ −*Z*_*b*_)/*Z*_*w*_, where *Z*_*w*_ and *Z*_*b*_ denote the mean within-network and between-network connectivity over the seven networks.

The remaining two markers were derived from task-evoked responses, estimated with a general linear model fitted to each run by ordinary least squares: the regressors were the boxcar time courses of the two conditions of the task (remembered and forgotten scenes in Scenes; hard and easy blocks in Words) convolved with the canonical double-gamma hemodynamic response function, together with an intercept and linear and quadratic drift terms, and the contrast between the two condition β values was taken in each of the 140 ROIs. The subsequent memory effect, the difference in encoding activity between later remembered and forgotten items, diminishes in older adults, including the loss of medial parietal deactivation^38,39^; memory encoding was therefore quantified by the expression of the subsequent memory effect, which measures how strongly the canonical spatial pattern of this effect is expressed in a run. It was computed from the Scenes runs by projecting the run’s map of the contrast *β*_*remembered*_− *β*_*forgotten*_ over the 140 ROIs, obtained by this model, onto the group-mean map of the observed data normalized to unit length. The modulation of activity by processing demand narrows with age^4,40,41^; demand adaptation was therefore quantified by demand modulation, which measures how strongly activity across the brain changes between the hard and easy conditions, regardless of the direction of the change. It was computed from the Words run as the contrast *β*_*hard*_− *β*_*easy*_ of the same model, taken in absolute value and averaged over the 140 ROIs.

### Model architecture

Brain-VRNN is a connectome-constrained variational recurrent neural network (V-RNN) that generates the BOLD of all four tasks in a top-down manner from three levels of latent states (Fig. 1a), following our previous model of macaque ECoG^29^ and the predictive-coding-inspired variational RNN^27,28,66^. The regional level, the lowest of the three, consists of 52 modules, one per brain region, each of which holds deterministic recurrent states d and probabilistic latent states z and outputs the BOLD of the ROIs that the region covers. The contextual level above it is a single module with slower dynamics that receives the global latent and sends its deterministic states to all 52 regional modules. The global level at the top is a two-dimensional latent that has no dynamics of its own and remains constant within a run. In the regional module k (k = 1, …, 52), the internal state 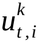 of unit i at time step t follows

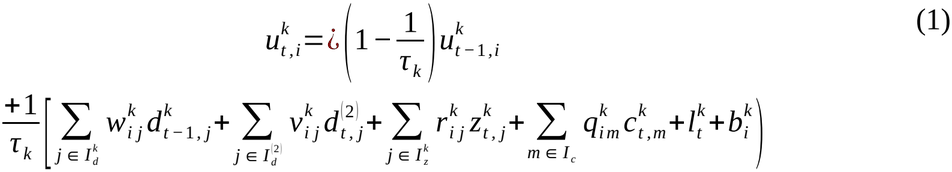

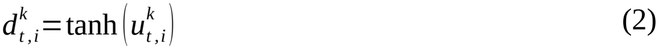

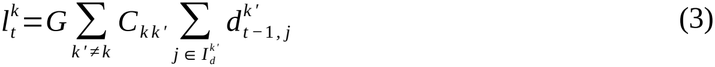

Here, the superscript k indexes the regional module (level 1) and the superscripts (2) and (3) denote the contextual and global levels; 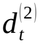 denotes the deterministic states of the contextual module, 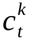 the condition vector of region k, *τ* _*k*_ the time constant, and *w, v, r, q*, and *b* the learned weights and biases. 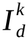 and 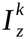 denote the index sets of the deterministic units and of the latent dimensions of regional module k, 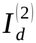 the index set of the deterministic units of the contextual module, and *I* _*c*_ the index set of the six elements of the condition vector; j and m index the elements of these sets, so that 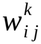, for example, is the weight from unit j to unit i. The lateral input 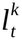 in Eq. (3) carries the activity of every other region k′ to region k in proportion to the structural connectivity *C*_*k k ′*_ of the person being modeled; the connectivity is fixed, and only the global coupling parameter G is learned^48,67^. The latent states are Gaussian with diagonal covariance, and their prior is parameterized by the deterministic states of the same module at the previous step,

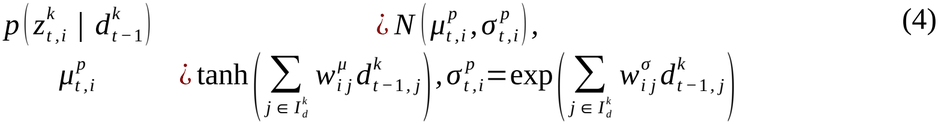

whereas the posterior is parameterized by adaptive variables *a*_*t*_ specific to each run,

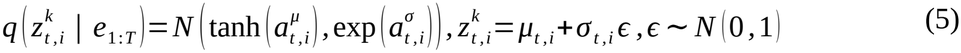

Here, *e*_1 : *T*_ denotes the prediction errors of the run, through which the adaptive variables are updated by backpropagation through time. The BOLD of ROI n covered by region k is read out from its deterministic states,

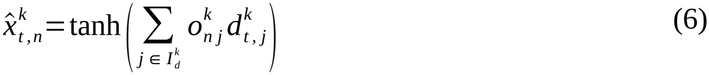

Here, 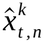 denotes the generated BOLD of ROI n at time step t, and *o*^*k*^ the learned readout weights of region k. The contextual module follows Eqs. (1), (2), (4), and (5) with the global latent 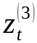 in place of 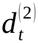 as its top-down input, without the lateral input 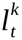, and with its own condition vector. The global latent *z*^(3)^ has a fixed prior *N* (0, 1) and a posterior parameterized by adaptive variables at the first step only, which are carried unchanged through the run, 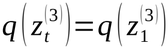 one dimension is designated trait and the other state (Eq. (7)).

The global latent was given two dimensions, the minimum for separating the trait-like from the state-like part of individuality, which was the aim of this study. Each regional module had 15 deterministic units, 2 latent dimensions, and a time constant of 2, and the contextual module had 15 deterministic units, 5 latent dimensions, and a time constant of 4. These sizes were chosen in preliminary experiments as the smallest configuration that reconstructed the training data stably: reducing the regional latent to one dimension worsened the reconstruction, whereas doubling the deterministic units or the contextual latent dimensions did not improve it. The time constants were taken from our previous model, in which larger values impaired the prediction of fast components^29^. The deterministic states of the regional and contextual modules therefore evolve on two time scales^31^, and the global latent sets run-level characteristics without evolving.

### Trait and state dimensions of the global latent

Let g(s) denote the group of runs with which run s shares its trait value; the group was either the person or the session. Let 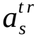 and 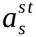 denote the adaptive variables (*a*^*μ*^, *a*^*σ*^) of the first and second dimensions of the global latent of run s, which are constant within a run. The first dimension, trait, took a single value per group, whereas the second dimension, state, was specific to each run. The constraint was enforced by projection: after every update of F, the trait variables of every run were replaced by the mean of the current values over the runs of its group,

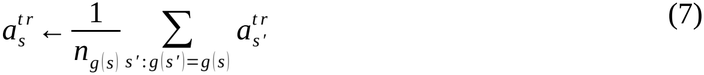

where *n*_*g*(*s*)_ is the number of runs of the group; this replacement is the Euclidean projection onto the set of runs with equal trait values and leaves 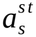 unchanged. During training, the group was the person, so that the trait dimension took one value per person across all tasks and all time points and any change between time points had to be carried by the state dimension. During latent-state estimation for unseen individuals, the group was the session, so that the trait dimension took a single value only among the tasks of one time point and no information was shared across time points. The stability of the estimated trait dimension across time points (Fig. 3a) is therefore not imposed by Eq. (7) but has to be recovered by the model from the BOLD of each time point alone.

### Training

The weights and the latent states of all training runs were learned jointly by minimizing the variational free energy^27,28,68^ summed over runs and time steps,

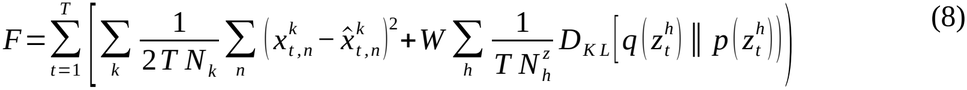

Here, h indexes the 54 modules, that is, the 52 regional modules, the contextual module, and the global module, 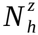 denotes the number of latent dimensions of module h, and *D*_*KL*_ the Kullback–Leibler divergence of the posterior q from the prior p. 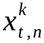 denotes the observed BOLD of ROI n at time step t, and the sum over n ranges over the *N*_*k*_ ROIs that region k covers. For the global module, whose latent is constant within a run, the divergence is evaluated once per run. The first term is the reconstruction error of the BOLD and the second term regularizes the posterior toward the prior; the meta-prior W balances the two and was set to the same value for all modules so as not to bias the flow of prediction error between levels^28^. Its value, 0.05, was chosen in preliminary experiments on an earlier version of the model over 0.005 to 0.1: a larger value (0.1) collapsed the amplitude of the generated BOLD, whereas a smaller value (0.02) degraded generation from the prior and the direction of the counterfactual response.

Let *θ* denote the parameters shared by all runs, that is, the weights *w*^*k*^, *v*^*k*^, *r*^*k*^, *q*^*k*^, *o*^*k*^, the biases *b*^*k*^, and the prior weights *w*^*μ*^, *w*^*σ*^ of every module, together with the global coupling G. The adaptive variables 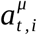 and 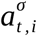 at all three levels are not part of *θ* but are specific to each run, and *a*_*s*_ denotes those of run s. Training minimized F summed over all training runs with respect to *θ* and to *a*_*s*_ of every training run jointly, with the structural connectivity *C*_*s*_ and the condition vector *c*_*s*_ of each run held fixed as inputs and the lateral weights held at *C*_*s*_. The weights were initialized by the Xavier method^69^, the biases were drawn from *N* (0, 10)^70^, and the global coupling parameter G was initialized at zero. *θ* and *a*_*s*_ of every training run were updated for 20,000 epochs on the full training set with the Rectified Adam optimizer^71^ (learning rate 5 × 10^−3^, reduced tenfold when the loss plateaued for 800 epochs, to a floor of 5 × 10^−4^; β_1_ = 0.9, β_2_ = 0.999) and a decoupled weight decay of 10^−4^ on the weights; the learning rate for G was one tenth of that for the weights. The gradient of F was propagated through the lateral pathway of Eq. (3) as well as through the recurrent and top-down connections, so that each region learned its dynamics under the influence it receives from the others; the lateral weights *C*_*k k ′*_ themselves were not updated, and the only parameter learned in this pathway was G. The learned global coupling converged to 0.008-0.027.

### Latent-state estimation for unseen individuals

After training, *θ* was frozen, and for each run s of a test individual only the run-specific variables were estimated from the observed BOLD: the adaptive variables *a*_*s*_ at all three levels, and an adjusted connectivity 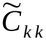 that replaces *C*_*k k ′*_ in the lateral input of Eq. (3) for that run^27,29^. This estimation is a form of data assimilation, in which the run-specific variables are fitted so that the model, with its learned dynamics, reproduces the observed BOLD of that individual. The estimation minimized F of Eq. (8) over the whole run with respect to these variables, given that person’s structural connectivity and condition values, using Rectified Adam (learning rate 10^−3^) for 20,000 iterations. The adaptive variables of the regional and contextual modules were initialized from the prior, and those of the global latent from the median over training runs. The adjusted connectivity was initialized at the measured *C*_*k k ′*_ and kept close to it by a quadratic penalty 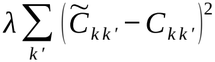 added to F (λ = 1, the value in a preliminary sweep at which the connectivity stayed anchored to the measured structure without raising the reconstruction error). Consequently, the estimated 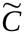 departed only slightly from the measured *C* (run-averaged r > 0.98). The projection of Eq. (7) was applied after every iteration with the session as the group.

Given the estimated latent states, BOLD was generated by running the model forward. In posterior generation, all latent states were sampled from their posteriors; in prior generation, the regional latent states were instead sampled from their priors of Eq. (4), while the contextual and global states remained at their posteriors, so that the regional dynamics were driven by the condition, top-down, and lateral inputs alone. Posterior generation reproduced the observed BOLD more closely and preserved the individual characteristics of the run more faithfully, whereas prior generation tended to yield larger effects of a condition shift. Accordingly, posterior generation was mainly used for the reproduction of BOLD, the analyses of the trait and state dimensions, and the counterfactual aging and intervention experiments, and prior generation for the focal aging and focal sparing experiments.

### Counterfactual aging and intervention

A counterfactual aging was generated by shifting the age element of the condition vector of every module while keeping everything else of the run fixed,

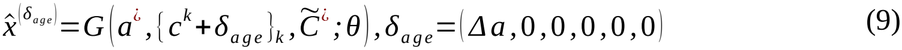

where *G* denotes forward generation by the model of Eqs. (1) to (6), *a*^¿^ the adaptive variables estimated for the run, 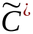 its adjusted connectivity, *θ* the frozen parameters, and δ_*age*_ a shift vector in the order of the elements of *c*_*t*_, whose age element *Δa* is given in units of the SD of age in the training set; *Δa*=0 gives the baseline generation. Each functional aging marker, written *m* (⋅) as a function of the BOLD from which it is computed, was recomputed from 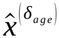. Age was shifted by *Δa* = −2, −1, +1, +2 (1 SD = 14.8 years), and the aging-related difference in the observed data was expressed per 1 SD of age for comparison. The sensitivity of each marker to age was defined for each session as the change per 1 SD of age,

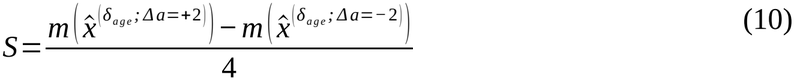

Preservation of individuality under counterfactual aging was checked by repeating the identification analysis described below at every dose. An intervention was generated by shifting, at the same time as age, the BMI element of the condition vector,

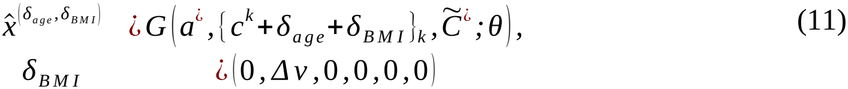

so that the two shifts act on different elements of the condition vector, 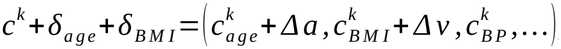, and 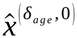 coincides with 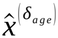 of Eq. (9).

Each run was regenerated under aging of *Δa*=+1 (+15 years) alone and under the same aging combined with a reduction of BMI by 3.0 kg/m^2^ (*Δv*=− 0.91); reductions of systolic blood pressure by 15 mmHg (−0.83 SD) and of amyloid burden by 0.9 SD were examined at comparable doses in the same form (Supplementary Fig. 1). For blood pressure, δ_*BP*_=(0, 0, *Δv*, 0, 0, 0) replaced δ_*BMI*_ in every module; for amyloid, δ_*amy*_=(0, 0, 0, 0, 0, *Δv*) was applied to the global burden supplied to the contextual module alone, leaving the regional deviations, that is, the spatial pattern of amyloid, unchanged. These doses were set at about 1 SD so that the shifted values stayed within a realistic range for this cohort. The decline D under aging alone and the recovery R produced by the intervention were defined as

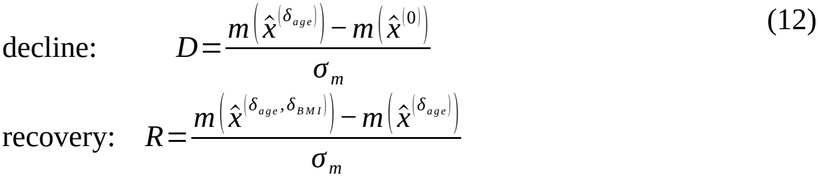

where *Δa*=+1 in both, *σ*_*m*_ is the SD of the marker in the observed data, and the fraction of the decline reversed is − *R* / *D*.

### Focal aging and focal sparing

The two experiments follow the same procedure (Fig. 5a) and differ only in which modules are aged and which generation serves as the baseline. For each of the 52 regions R in turn:

1. **Condition**. In focal aging, the age element of the condition vector was set to +2 SD for region R only, and the other 51 regions and the contextual module stayed at the person’s actual age. In focal sparing, region R stayed at the actual age, and the other 51 regions and the contextual module were set to +2 SD.
2. **Generation**. The BOLD of each run was regenerated by prior generation, with the estimated latent states and the adjusted connectivity kept fixed.
3. **Markers**. The four functional aging markers were computed from the regenerated BOLD.
4. **Effect**. The markers of the baseline generation were subtracted: for focal aging, the baseline was the generation in which no region was aged; for focal sparing, the generation in which every region was aged. In both cases the two generations differ only in region R. The effects are denoted *Δ*_*aging*_ (*R*) and *Δ*_*sparing*_ (*R*).

Both effects were computed for each session, averaged over the sessions of each fold, and averaged over the two folds; homologous regions of the two hemispheres were combined, giving 26 values per marker. The correspondence between the two experiments was quantified as the Spearman correlation between *Δ*_*aging*_ and *Δ*_*sparing*_ over the 26 regions of the four markers combined, and as the overlap between the five regions with the largest effects in each experiment, counted per marker. Comparisons of regional maps between markers used the centered effects and are reported as correlations, which do not depend on the choice of baseline. Reproducibility of each map was assessed by split-half: the test individuals were divided at random into two halves, the maps computed from the two halves were correlated, and the median over 200 random divisions was adjusted by the Spearman–Brown formula.

### Evaluation of generated BOLD

Reproduction of the time course was quantified as the Pearson correlation between the observed and generated BOLD of each ROI in each test run, summarized as the median and IQR over ROIs and runs. Reproduction of the regional distribution of task responses was quantified by correlating, across the 140 ROIs, the group-mean maps of the subsequent memory contrast and of the demand contrast computed from the generated and from the observed BOLD. Individual specificity was assessed with resting-state functional connectivity among the 52 regions, computed from time series averaged over the ROIs of each region: for each test participant, the connectivity computed from the generated BOLD of all resting-state runs was correlated with the observed connectivity of the same participant and with that of every other test participant, and the participant was identified as the one with the highest correlation (n = 143 and 147 participants with resting-state runs in the two folds; chance level 1/143 and 1/147, about 0.007); the similarities and the accuracy were averaged over the two folds. The functional aging markers computed from the generated BOLD were compared with those from the observed BOLD by correlation across individuals and, within individuals, across time points. Each cognitive score was related to the functional aging markers in two ways: by the Pearson correlation with each marker separately (Supplementary Tables 1 and 2), and by the standardized coefficients of a linear regression on the four markers entered together, which give the unique contribution of each marker (Supplementary Tables 3 and 4). The sessions of the two test halves were combined into a single sample, so that every participant was represented once by the model that had not seen them. These relations were compared between generated and observed BOLD over the 16 tests and PC1, as the correlation between the two sets of coefficients.

### Analysis of the trait and state dimensions

The trait and state dimensions were examined at test, where no tying was applied across time points. The sign of each dimension is arbitrary in each fold, so the orientation was fixed by the convention that the correlation of its values with sex (coded 1 for women) is positive; higher and lower values refer to this orientation throughout. The convention is independent of the variables used in the analyses, and neither fold required a flip. Before the two folds were pooled, the trait and state values were z-scored within each fold.

Stability across time points was quantified for individuals with two or more sessions as the Pearson correlation between the values at two time points: within each task, values were averaged over the runs of a session and z-scored within each fold, task, and pair of time points, and all pairs were pooled into a single correlation. The intraclass correlation coefficient (ICC)^72^ was computed as the between-person variance divided by the sum of the between- and within-person variances, with values z-scored within each fold and pooled across folds; because the trait dimension is shared within a session, the ICC of its values was computed on session values, whereas that of the state dimension, which varies from run to run, was computed on run values.

Relations to external variables were quantified as semipartial correlations at the session level, with the value of the other dimension partialled out. The prediction of future cognition was tested with the model *score*_*b*_ ~ *score*_*a*_ +*trait* fitted over all transitions between consecutive time points stacked together, with the trait value estimated from the BOLD of the earlier time point only and Benjamini–Hochberg correction^73^ over the 12 longitudinal measures; the same model was fitted with the state value in place of the trait value.

The relation of the age sensitivity to the trait value was tested with the model |*S*) ~ *baseline* +*age* +*trait* + *state* at the session level, where baseline is the marker generated at the actual condition values. The decline and the recovery in the intervention experiment were modeled in the same form, and the ratio between individuals with low and high trait values was obtained from the predicted values at trait values of −1 and +1 SD; the possibility of a ceiling effect was examined by adding nonlinear terms of baseline, which attenuated the effect of the trait value by at most 16%. In all of these regressions, standard errors were clustered by participant.

### Statistical analysis

Confidence intervals and the p-values for the difference between the two dimensions were obtained by bootstrap over participants (2000 to 4000 resamples). All tests were two-tailed, and confidence intervals are 95% intervals.

### Use of large language models

Large language model-based assistants were used to edit the language of the text and to assist in writing and checking analysis code; the authors reviewed all output and take full responsibility for the content.

## Data availability

The DLBS data are available from OpenNeuro (ds004856) at https://openneuro.org/datasets/ds004856^33^.

## Code availability

The model was implemented in C++ with MPI and the analyses in Python. The code will be made publicly available upon formal publication of this article.

## ACKNOWLEDGEMENTS

Data were provided by the Dallas Lifespan Brain Study (OpenNeuro ds004856); we thank the DLBS investigators and participants for making the data available. This work was supported by the JSPS KAKENHI (JP21K15723, JP24K20897, JP24H00076, JP24K00499, JP25H01173), JST BOOST (JPMJBY24E5), JST ACT-X (JPMJAX24C2), JST CREST (JPMJCR21P4), AMED (JP25wm0625419, JP21tm0424601, JP24wm0625407), JH Research Fund (JH2026-A-01), and Intramural Research Grant (6-9, 7-9, 8-1) for Neurological and Psychiatric Disorders of NCNP. The funders played no role in the study design, data collection, analysis, data interpretation, or manuscript writing.

## AUTHOR CONTRIBUTIONS

Y.T. and Y.Y. conceived the study and designed its core framework. Y.T. conducted the experiments and analysis and prepared the original draft under the supervision of Y.Y. H.I. developed the original model code and contributed to the architecture and experimental protocols, and H.T. built the computational environment. All authors reviewed the manuscript, contributed to its revision, and approved the final version.

## COMPETING INTERESTS

The authors declare no competing interests.

## Notes

### Competing Interest Statement

The authors have declared no competing interest.

