## Supplementary Information for "Counterfactual aging in individual digital twin brains uncovers a latent dimension of resilience to aging"

**Supplementary Figure 1:** Blood pressure and amyloid burden as interventions at a dose matched to BMI.

**Supplementary Table 1:** Correlations between the four functional aging markers computed from the observed BOLD and the cognitive test scores.

**Supplementary Table 2:** Correlations between the four functional aging markers computed from the generated BOLD and the cognitive test scores.

**Supplementary Table 3:** Unique contributions of the four functional aging markers computed from the observed BOLD to the cognitive test scores.

**Supplementary Table 4:** Unique contributions of the four functional aging markers computed from the generated BOLD to the cognitive test scores.

**Supplementary Figure 1. Blood pressure and amyloid burden as interventions at a dose matched to BMI.**

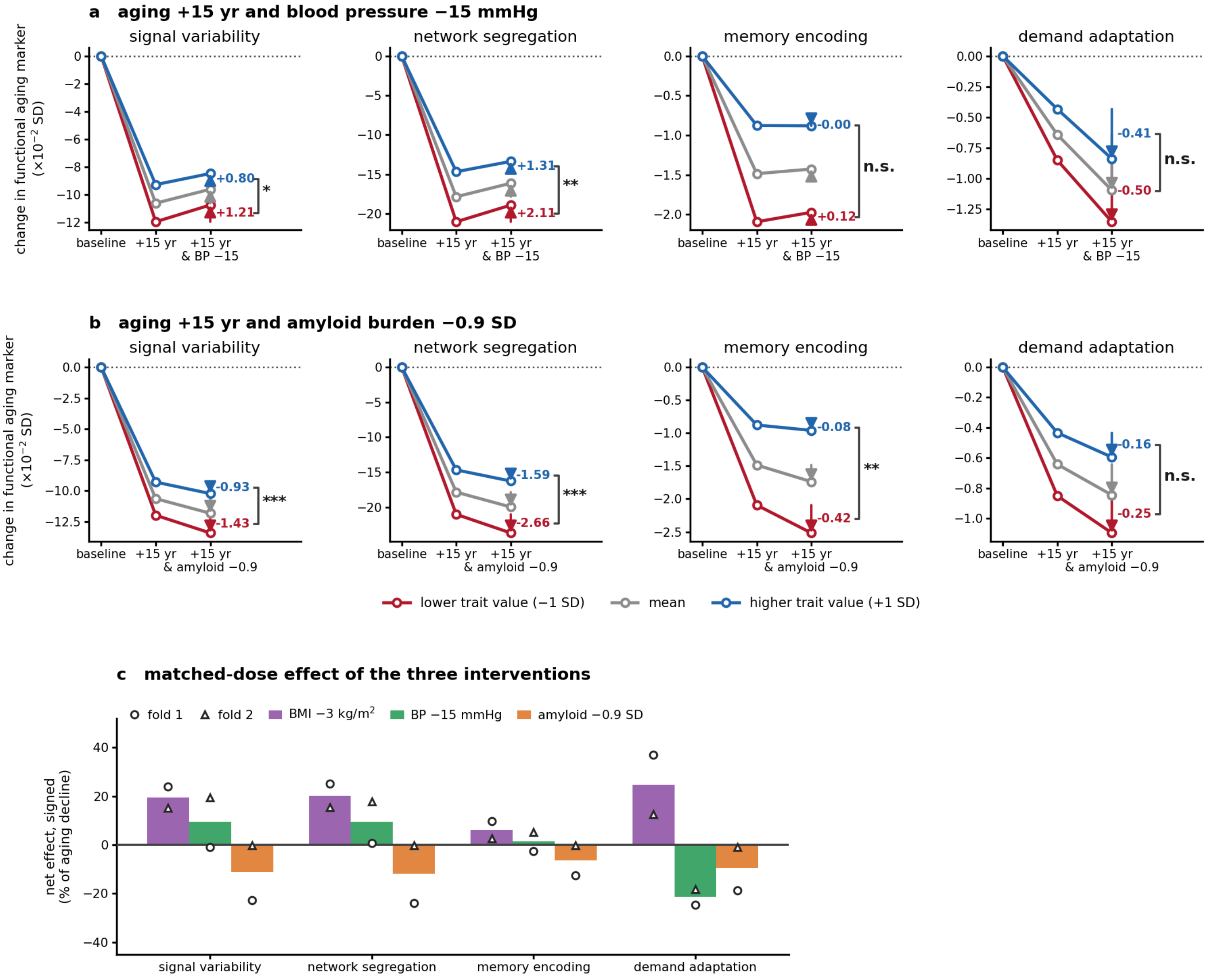


**a**, Change in each functional aging marker (×10$^{-2}$ SD) relative to the baseline generation at each individual's actual age, under aging alone (+15 years, +1 SD) and under the same aging combined with a reduction of systolic blood pressure by 15 mmHg (−0.83 SD), in the same form as Fig. 6c, for individuals with a lower (1 SD below the mean; red), average (gray), and higher (1 SD above the mean; blue) trait value. Numbers, the intervention step at trait values of −1 and +1 SD; asterisks, effect of the trait value on the step (cluster-robust regression pooled over the two folds; *, $p < 0.05$; **, $p < 0.01$; ***, $p < 0.001$; n.s., not significant). **b**, The same for a reduction of amyloid burden by 0.9 SD. **c**, Net effect of the three interventions on each marker, expressed as a percentage of the aging change: positive, the marker moves back toward its baseline value; negative, further in the direction of aging. All three conditions are shifted by about −0.9 SD of their own distribution (BMI −0.91 SD; systolic blood pressure −0.83 SD; amyloid burden −0.9 SD) on the identical aging arm. Bars, the two folds pooled; open symbols, each fold separately (circle, fold 1; triangle, fold 2). BMI consistently moved the markers back toward their baseline values, whereas blood pressure and amyloid burden did not move them consistently in that direction and disagreed between the folds. BMI, body mass index; BP, systolic blood pressure.

**Supplementary Table 1. Correlations between the four functional aging markers computed from the observed BOLD and the cognitive test scores.**

| Test | Domain | n | Signal variability | Network segregation | Memory encoding | Demand adaptation |
|---|---|---|---|---|---|---|
| MMSE | Screening | 532 | +0.09 | +0.13* | +0.13* | +0.14* |
| Digit comparison | Processing speed | 532 | +0.35* | +0.33* | +0.09 | +0.34* |
| Digit symbol | Processing speed | 532 | +0.38* | +0.35* | +0.14* | +0.39* |
| Spatial working memory (errors) | Working memory | 532 | -0.30* | -0.19* | -0.10* | -0.27* |
| Letter–number sequencing | Working memory | 532 | +0.29* | +0.25* | +0.08 | +0.27* |
| Delayed matching to sample (CANTAB) | Working memory | 260 | +0.32* | +0.25* | +0.12 | +0.25* |
| Spatial recognition memory (CANTAB) | Working memory | 260 | +0.11 | +0.16* | +0.06 | +0.08 |
| HVLT delayed recognition | Episodic memory | 532 | +0.17* | +0.11* | +0.08 | +0.17* |
| Woodcock–Johnson delayed recall | Episodic memory | 442 | +0.41* | +0.32* | +0.21* | +0.42* |
| Verbal recognition memory (CANTAB) | Episodic memory | 259 | +0.23* | +0.21* | +0.05 | +0.17* |
| Raven matrices | Reasoning | 532 | +0.32* | +0.18* | +0.20* | +0.32* |
| Letter sets | Reasoning | 532 | +0.34* | +0.18* | +0.08 | +0.36* |
| Stockings of Cambridge (CANTAB) | Reasoning | 532 | +0.20* | +0.12* | +0.15* | +0.24* |
| ETS vocabulary | Vocabulary | 532 | +0.00 | -0.09 | -0.02 | +0.14* |
| Graded naming | Vocabulary | 428 | +0.05 | -0.04 | +0.06 | +0.15* |
| Controlled oral word association | Verbal fluency | 260 | +0.17* | +0.21* | +0.01 | +0.28* |
| General cognitive score (PC1) | Composite | 532 | +0.41* | +0.30* | +0.17* | +0.43* |

Pearson correlation (r) between each functional aging marker and each cognitive score across sessions, with the sessions of a participant at different time points treated as separate observations, pooled over the test halves of the two folds (up to 294 participants); n, number of sessions with the score available. Spatial working memory is scored by errors, so a negative coefficient indicates better performance at a higher value of the marker. PC1, general cognitive score (Methods). Asterisks, $q < 0.05$ after Benjamini–Hochberg correction over the 68 correlations of the table (4 markers × 17 scores).

**Supplementary Table 2. Correlations between the four functional aging markers computed from the generated BOLD and the cognitive test scores.**

| Test | Domain | n | Signal variability | Network segregation | Memory encoding | Demand adaptation |
|---|---|---|---|---|---|---|
| MMSE | Screening | 532 | +0.11* | +0.14* | +0.14* | +0.17* |
| Digit comparison | Processing speed | 532 | +0.37* | +0.31* | +0.08 | +0.34* |
| Digit symbol | Processing speed | 532 | +0.39* | +0.34* | +0.15* | +0.39* |
| Spatial working memory (errors) | Working memory | 532 | -0.28* | -0.15* | -0.08 | -0.24* |
| Letter–number sequencing | Working memory | 532 | +0.27* | +0.20* | +0.06 | +0.26* |
| Delayed matching to sample (CANTAB) | Working memory | 260 | +0.29* | +0.19* | +0.11 | +0.22* |
| Spatial recognition memory (CANTAB) | Working memory | 260 | +0.12 | +0.13* | +0.03 | +0.10 |
| HVLT delayed recognition | Episodic memory | 532 | +0.14* | +0.10* | +0.09 | +0.15* |
| Woodcock–Johnson delayed recall | Episodic memory | 442 | +0.41* | +0.29* | +0.20* | +0.39* |
| Verbal recognition memory (CANTAB) | Episodic memory | 259 | +0.22* | +0.19* | +0.00 | +0.16* |
| Raven matrices | Reasoning | 532 | +0.31* | +0.15* | +0.18* | +0.30* |
| Letter sets | Reasoning | 532 | +0.33* | +0.17* | +0.07 | +0.33* |
| Stockings of Cambridge (CANTAB) | Reasoning | 532 | +0.16* | +0.08 | +0.16* | +0.22* |
| ETS vocabulary | Vocabulary | 532 | -0.02 | -0.08 | -0.04 | +0.14* |
| Graded naming | Vocabulary | 428 | +0.02 | -0.02 | +0.05 | +0.15* |
| Controlled oral word association | Verbal fluency | 260 | +0.20* | +0.22* | +0.01 | +0.29* |
| General cognitive score (PC1) | Composite | 532 | +0.40* | +0.27* | +0.16* | +0.41* |

As in Supplementary Table 1, with the four markers computed from the BOLD generated from the estimated latent states of the same sessions (posterior generation without intervention) instead of the observed BOLD. The 68 coefficients correlate with those of Supplementary Table 1 at r = 0.99.

**Supplementary Table 3. Unique contributions of the four functional aging markers computed from the observed BOLD to the cognitive test scores.**

| Test | Domain | n | Signal variability | Network segregation | Memory encoding | Demand adaptation | $R^2$ |
|---|---|---|---|---|---|---|---|
| MMSE | Screening | 427 | +0.01 | +0.03 | +0.11 | +0.11 | 0.03 |
| Digit comparison | Processing speed | 427 | +0.10 | +0.20* | +0.02 | +0.22* | 0.17 |
| Digit symbol | Processing speed | 427 | +0.12 | +0.21* | +0.06 | +0.27* | 0.23 |
| Spatial working memory (errors) | Working memory | 427 | -0.21* | -0.06 | -0.03 | -0.12 | 0.11 |
| Letter–number sequencing | Working memory | 427 | +0.18* | +0.11 | +0.02 | +0.13* | 0.12 |
| Delayed matching to sample (CANTAB) | Working memory | 213 | +0.17 | +0.09 | +0.09 | +0.13 | 0.11 |
| Spatial recognition memory (CANTAB) | Working memory | 213 | +0.06 | +0.13 | +0.04 | +0.03 | 0.04 |
| HVLT delayed recognition | Episodic memory | 427 | +0.09 | +0.05 | +0.05 | +0.14* | 0.06 |
| Woodcock–Johnson delayed recall | Episodic memory | 358 | +0.15* | +0.16* | +0.12* | +0.28* | 0.25 |
| Verbal recognition memory (CANTAB) | Episodic memory | 212 | +0.18 | +0.11 | +0.00 | +0.03 | 0.07 |
| Raven matrices | Reasoning | 427 | +0.17* | -0.01 | +0.15* | +0.22* | 0.15 |
| Letter sets | Reasoning | 427 | +0.24* | -0.02 | +0.01 | +0.24* | 0.17 |
| Stockings of Cambridge (CANTAB) | Reasoning | 427 | +0.08 | +0.04 | +0.11 | +0.18* | 0.08 |
| ETS vocabulary | Vocabulary | 427 | +0.01 | -0.17* | +0.00 | +0.19* | 0.05 |
| Graded naming | Vocabulary | 343 | +0.06 | -0.11 | +0.06 | +0.12 | 0.03 |
| Controlled oral word association | Verbal fluency | 212 | -0.04 | +0.12 | +0.00 | +0.25* | 0.08 |
| General cognitive score (PC1) | Composite | 427 | +0.21* | +0.09 | +0.09 | +0.28* | 0.24 |

Standardized coefficients (β) from a linear regression of each z-scored cognitive score on the four z-scored functional aging markers entered together, across sessions pooled over the test halves of the two folds (up to 267 participants); n, number of sessions with the score and all four markers available. $R^2$, proportion of variance explained by the four markers. Asterisks, $q < 0.05$ after Benjamini–Hochberg correction over the 68 coefficients of the table (4 markers × 17 scores).

**Supplementary Table 4. Unique contributions of the four functional aging markers computed from the generated BOLD to the cognitive test scores.**

| Test | Domain | n | Signal variability | Network segregation | Memory encoding | Demand adaptation | $R^2$ |
|---|---|---|---|---|---|---|---|
| MMSE | Screening | 427 | +0.03 | +0.02 | +0.12* | +0.14* | 0.04 |
| Digit comparison | Processing speed | 427 | +0.16* | +0.16* | +0.01 | +0.22* | 0.18 |
| Digit symbol | Processing speed | 427 | +0.17* | +0.17* | +0.08 | +0.26* | 0.24 |
| Spatial working memory (errors) | Working memory | 427 | -0.22* | -0.02 | -0.03 | -0.12 | 0.10 |
| Letter–number sequencing | Working memory | 427 | +0.18* | +0.05 | +0.02 | +0.15* | 0.10 |
| Delayed matching to sample (CANTAB) | Working memory | 213 | +0.21* | +0.01 | +0.09 | +0.11 | 0.09 |
| Spatial recognition memory (CANTAB) | Working memory | 213 | +0.08 | +0.10 | +0.02 | +0.03 | 0.03 |
| HVLT delayed recognition | Episodic memory | 427 | +0.07 | +0.04 | +0.08 | +0.14* | 0.05 |
| Woodcock–Johnson delayed recall | Episodic memory | 358 | +0.22* | +0.11 | +0.12* | +0.26* | 0.25 |
| Verbal recognition memory (CANTAB) | Episodic memory | 212 | +0.19 | +0.08 | -0.02 | +0.04 | 0.07 |
| Raven matrices | Reasoning | 427 | +0.21* | -0.05 | +0.15* | +0.20* | 0.14 |
| Letter sets | Reasoning | 427 | +0.27* | -0.06 | +0.01 | +0.23* | 0.16 |
| Stockings of Cambridge (CANTAB) | Reasoning | 427 | +0.09 | -0.01 | +0.12* | +0.17* | 0.07 |
| ETS vocabulary | Vocabulary | 427 | +0.00 | -0.15* | -0.03 | +0.19* | 0.04 |
| Graded naming | Vocabulary | 343 | +0.03 | -0.07 | +0.06 | +0.12 | 0.02 |
| Controlled oral word association | Verbal fluency | 212 | -0.00 | +0.12 | +0.01 | +0.24* | 0.09 |
| General cognitive score (PC1) | Composite | 427 | +0.24* | +0.04 | +0.10 | +0.28* | 0.23 |

As in Supplementary Table 3, with the four markers computed from the generated BOLD of the same sessions. The 68 coefficients correlate with those of Supplementary Table 3 at r = 0.96.
